# Using the NCBI AMRFinder Tool to Determine Antimicrobial Resistance Genotype-Phenotype Correlations Within a Collection of NARMS Isolates

**DOI:** 10.1101/550707

**Authors:** Michael Feldgarden, Vyacheslav Brover, Daniel H. Haft, Arjun B. Prasad, Douglas J. Slotta, Igor Tolstoy, Gregory H. Tyson, Shaohua Zhao, Chih-Hao Hsu, Patrick F. McDermott, Daniel A. Tadesse, Cesar Morales, Mustafa Simmons, Glenn Tillman, Jamie Wasilenko, Jason P. Folster, William Klimke

## Abstract

Antimicrobial resistance (AMR) is a major public health problem that requires publicly available tools for rapid analysis. To identify acquired AMR genes in whole genome sequences, the National Center for Biotechnology Information (NCBI) has produced a high-quality, curated, AMR gene reference database consisting of up-to-date protein and gene nomenclature, a set of hidden Markov models (HMMs), and a curated protein family hierarchy. Currently, the Bacterial Antimicrobial Resistance Reference Gene Database contains 4,579 antimicrobial resistance gene proteins and more than 560 HMMs.

Here, we describe AMRFinder, a tool that uses this reference dataset to identify AMR genes. To assess the predictive ability of AMRFinder, we measured the consistency between predicted AMR genotypes from AMRFinder against resistance phenotypes of 6,242 isolates from the National Antimicrobial Resistance Monitoring System (NARMS). This included 5,425 *Salmonella enterica*, 770 *Campylobacter* spp., and 47 *Escherichia coli* phenotypically tested against various antimicrobial agents. Of 87,679 susceptibility tests performed, 98.4% were consistent with predictions.

To assess the accuracy of AMRFinder, we compared its gene symbol output with that of a 2017 version of ResFinder, another publicly available resistance gene database. Most gene calls were identical, but there were 1,229 gene symbol differences between them, with differences due to both algorithmic differences and database composition. AMRFinder missed 16 loci that Resfinder found, while Resfinder missed 1,147 loci AMRFinder identified. Two missing drug classes from the 2017 version of ResFinder contributed 81% of missed loci. Based on these results, AMRFinder appears to be a highly accurate AMR gene detection system.

**Importance:** Antimicrobial resistance is a major public health problem. Traditionally, antimicrobial resistance has been identified using phenotypic assays. With the advent of genome sequencing, we now can identify resistance genes and deduce if an isolate could be resistant to antibiotics. We describe a database of 4,579 acquired antimicrobial resistance genes, the largest publicly available, and a software tool to identify genes in bacterial genomes, AMRFinder. Unlike other tools, AMRFinder uses a gene hierarchy to prevent overpredicting what the correct gene call should be, enabling more accurate assessment. To assess these resources, we determined the resistance gene content of over 6,200 bacterial isolates from the National Antimicrobial Resistance Monitoring System that have been assayed using traditional methods and that also have had their genomes sequenced. We also compared our gene assessments to those of a popularly used tool. We found that AMRFinder has a high overall consistency between genotypes and phenotypes.

## Introduction

Antimicrobial resistance (AMR) is a major public health problem, with an estimated 23,000 deaths annually in the U.S. attributable to antimicrobial resistant infections (https://www.cdc.gov/drugresistance/threat-report-2013/index.html). The continued evolution of multi-drug resistance ensures that AMR will continue to be a health challenge for years to come. As described in the National Strategy on Combating Antibiotic Resistant Bacteria report (https://www.cdc.gov/drugresistance/pdf/national_action_plan_for_combating_antibotic-resistant_bacteria.pdf), there is a critical need to understand how AMR is related to bacterial genotype, both to enhance AMR mechanism discovery and to enable AMR diagnostics. One key method to establish this link is genome sequencing, which can also be used for surveillance purposes.

Traditionally, AMR has been identified using phenotypic assays. The gold standard for measuring antimicrobial susceptibility is based on standardized dilution- or diffusion-based *in vitro* antimicrobial susceptibility testing (AST) methods, where extensive research and testing have been performed to correlate AST measurements with clinical outcomes (1) Increasingly, molecular methods are being used in resistance surveillance and in some cases also to guide clinical therapy. These range from PCR detection of known resistance elements (2) to mass spectrometry-based methods (3-7). Whole genome shotgun sequencing (WGS) has been integrated into the clinical and public health settings, though the use of WGS has focused primarily on outbreak identification and tracking (8,9). Along with epidemiological uses, there is great potential for the use of WGS to aid and guide AMR detection (10-15). Accurate assessment of AMR gene content enables the discovery of novel resistance variants and can serve as the basis for predicting resistance phenotypes without the need for time consuming phenotypic tests (11,16,17).

An in-silico approach to assessing AMR content requires comprehensive and accurate AMR gene databases as well as tools that can reliably identify AMR genes. There are many databases and tools using a variety of approaches and data sources as described in a recent review (18). While some tools exclusively use BLAST-based approaches (19), others incorporate Hidden Markov Model (HMM) approaches (20). BLAST-based approaches are able to identify specific alleles and closely-related genes. However, BLAST-based methods use arbitrary cutoffs that can miscall AMR genes or even misattribute resistance to non-AMR genes (e.g., misidentification of metallo-beta-hydrolases as metallo-beta-lactamases(21)). HMM approaches facilitate a hierarchical classification of AMR proteins, from alleles to gene families, but curation and validation of HMM libraries are required. Tools also differ based on whether they analyze nucleotide or protein sequence. Additionally, some tools are only available through a web-interface, while others can be operated on local servers providing more flexibility to users. Researchers attempting to use currently available AMR databases must choose between these different database resources. Some contain collections of alignments of resistance genes for use in HMMs (20). Others consist of collections of nucleotide or protein sequences of either individual resistance genes or resistance-related mobile elements (22,23). Some databases are actively curated such as the CARD (23,24), ResFinder (22), and the Lahey Clinic database (https://www.lahey.org/Studies/; the latter is now hosted and maintained by NCBI, as part of the NCBI’s Bacterial Antimicrobial Resistance Reference Gene Database), while others are not actively updated. Separate groups curate different classes of genes, and even a single class of genes can be curated by multiple groups (e.g., beta-lactamases). In addition, some data resources include allelic variation of housekeeping genes that can confer or contribute to resistance, while others focus exclusively on acquired resistance mechanisms. Assessing and comparing these resources and tools is also challenging as there are few high-quality strain collections that have been extensively genotyped and phenotyped, and that are also publicly available.

Here, we describe the development of a comprehensive AMR gene database, the Bacterial Antimicrobial Resistance Reference Gene Database, and the development of AMRFinder, an AMR gene identification tool, along with publicly available datasets to test AMR gene detection methods. To identify AMR genes from sequence data, we created over 560 AMR HMMs (21) and curated over 4,579 AMR protein sequences, placing both in a hierarchical framework of gene families, symbols, and names in collaboration with multiple groups including CARD (21). We then developed AMRFinder to leverage both the content and structure of this database to accurately identify and name AMR gene sequences. To validate this system, we used a collection of isolates from the NARMS program that have undergone extensive susceptibility testing and whole genome shotgun assembly, and we also compared AMRFinder performance with a version of ResFinder 2.0 released in 2017.

## Methods

### AMR gene database

The Bacterial Antimicrobial Resistance Reference Gene Database contains a hierarchy of AMR protein families and is stored in NCBI’s RefSeq database (21). Each protein, and each protein family, has a curated name and gene symbol where appropriate. Gene symbols can point to more than one protein sequence, while alleles point to one unique amino acid sequence. For many families, we have constructed protein HMMs that identify these protein families. When necessary, the protein sequence has been manually verified to be full-length and to have the appropriate start site. The proteins are arranged in protein family hierarchies based on protein homology and function.

Our collection of AMR proteins is derived from multiple sources, including the compilation of beta-lactamase alleles and Qnr family quinolone resistance protein alleles compiled by the Lahey Clinic team (http://www.lahey.org/studies/ (25), ResFinder (22), and the Comprehensive Antimicrobial Resistance Database [CARD; (24)]. At the request of the Lahey Clinic team of Drs. Karen Bush, George Jacoby, and Timothy Palzkill (https://www.lahey.org/Studies/), NCBI has assumed responsibility for assigning and curating beta-lactamase alleles (https://www.ncbi.nlm.nih.gov/pathogens/submit-beta-lactamase/). The assignment process uses many beta-lactamase subfamily HMMs that are also used by AMRFinder. Families covered include the 27 previously covered by Lahey, the ADC and PDC families, as well as the newly assigned families CMH, CRH, and FRI. Since January 2016, NCBI has assigned 676 new beta-lactamase alleles. These newly assigned alleles as well as those previously curated are incorporated into our AMR gene database. We obtained compilations of resistance genes for several classes of ribosome-targeting antibiotics from Dr. Marilyn Roberts [(26) and personal communication]. We obtained collections of AMR proteins encoded in integron regions from both RAC (27) and INTEGRALL (28). Additional sources included compilations provided by collaborating groups such as the FDA Center for Veterinary Medicine, University of Oxford (Dr. Derrick Crook), and the *Klebsiella* Sequence Typing Database at the Pasteur Institute (http://bigsdb.pasteur.fr/klebsiella/klebsiella.html). These sources were supplemented by continuous examination of review articles and new reports of resistance proteins.

The 4,528 resistance proteins in our database as of this writing confer resistance to 34 classes of antimicrobials and disinfectants, and are encoded by over 800 gene families. All underlying nucleotide records contain complete coding sequence and are not derived from synthetic constructs. Nucleotide sequences were oriented with the AMR protein coding region on the positive strand, and records were constructed, where possible, to include an additional 100bp on either side of the coding region to assist in the design of primers. Protein records were created as described previously (21). This collection has a standardized nomenclature to provide maximal functional information as well as ease of bioinformatic use, and is found under in our Reference Gene Browser (https://www.ncbi.nlm.nih.gov/pathogens/isolates#/refgene/) as well as RefSeq BioProject PRJNA313047 (https://www.ncbi.nlm.nih.gov/bioproject/PRJNA313047).

### AMR HMM construction

Groups of related AMR proteins with similar sequences and similar gene symbols as taken from our various sources were aligned using MUSCLE (29) or Clustal W (30), then viewed, trimmed, and culled of mis-assigned, redundant, frameshifted, or fragmentary sequences, using Belvu (31). The resulting curated multiple sequence “seed” alignments were used to construct protein profile HMMs, using the HMMER3 package (http://hmmer.org/). In some cases, BLAST or HMM searches recruited additional sequences that were judged valid to add to the seed alignments so that the scores obtained in HMM search results could more clearly separate true family members from outgroup sequences. The ResFams (20) library of HMMs, based on sequences taken from CARD sequences and clustered by their CARD antibiotic resistance ontology assignments, provided important early assistance in recognizing putative AMR proteins and grouping them into homology families. However, to create a hierarchical classification system for AMR proteins, with sufficiently fine divisions of recognized families and cutoffs values that could prove trustworthy while searching very large data sets, we created, calibrated, and annotated an entirely new HMM library, available at https://ftp.ncbi.nlm.nih.gov/hmm/NCBIfam-AMRFinder/. The literature was reviewed, molecular phylogenetic trees and search results were examined, and an informative protein name was selected for each HMM built to represent a family of AMR proteins. These HMMs support correct functional annotation of AMR proteins for RefSeq prokaryotic genomes (21).

### Identifying acquired AMR genes

#### Protein searches

AMRFinder-prot uses the database of AMR gene sequences, HMMs, the hierarchical tree of AMR protein designations, and a custom rule-set to generate names and coordinates for AMR genes, along with descriptions of the evidence used to identify the sequence. Software and documentation are available at https://github.com/ncbi/amr and https://www.ncbi.nlm.nih.gov/pathogens/antimicrobial-resistance/AMRFinder/. Genes are reported with the following procedure after both HMMER and BLASTP searches are run.

#### BLASTP matches

In AMRFinder, BLASTP (32,33) is run with the -task blastp-fast - word_size 6 -threshold 21 -evalue 1e-20 -comp_based_stats 0 options against the AMR gene database described above. Exact BLAST matches over the full length of the reference protein are reported. If there is no exact match, then the following rules are applied: Matches with < 90% identity or with < 50% coverage of the protein are dropped. If the hit is to a fusion protein then at least 90% of the protein must be covered. A BLAST match to a reference protein is removed if it is covered by another BLAST match which has more identical residues or the same number of identical residues, but to a longer reference protein. A single match is chosen as the best of what remains sorting by the following criteria in order (1) if it is exact; (2) has more identical residues; (3) hits a shorter protein; or (4) the gene symbol comes first in alphabetical order.

#### HMM matches

HMMER version 3.1b2 (http://hmmer.org/) is run using the --cut_tc -Z 10000 options with the HMM database described above. HMM matches with full_score < TC1 or domain_score < TC2 are dropped. All HMM matches to HMMs for parent nodes of other HMM matches in the hierarchy are removed. The match(es) with the highest full score are kept. If there is an exact BLAST match or the family of the BLAST match reference protein is descendant of the family of the HMM then the information for the nearest HMM node to the BLAST match are returned.

#### Translated DNA searches

Translated alignments using BLASTX of the assembly against the AMR protein database were used to help identify partial, split, or unannotated AMR proteins using the -task tblastn-fast -word_size 3 -evalue 1e-20 -seg no -comp_based_stats 0 options. The algorithm for selecting hits is as described above for proteins, but note that HMM searches are not performed against the unannotated assembly.

#### Nucleotide searches

Nucleotide-nucleotide BLAST searches were also performed for evaluation purposes, although this is not built into AMRFinder. We collected the nucleotide sequences for all proteins in GenBank with sequences identical to those in the AMR database. The genome assembly for each isolate was masked at locations identified as AMR genes by AMRFinder before aligning the remainder against the nucleotide sequences we collected above. Hits were combined to determine coverage of the reference protein and all 7 hits with > 50% length and > 90% sequence similarity to a reference sequence were selected for analysis.

### Samples

The 6,242 isolates used in this study are from various NARMS projects (34) including 294 *Campylobacter coli*, 476 *Campylobacter jejuni*, 47 *Escherichia coli*, and 5,425 *Salmonella enterica*. Sources for these isolates include human clinical *S. enterica* isolates resistant to at least one antibiotic from 2014, NARMS food animal cecal testing projects, food adulterant isolates including Shiga-toxin producing *E. coli*, and routine NARMS retail meat surveillance. Isolates are listed in Table S1 and are deposited in the Sequence Read Archive, or were independently assembled and submitted to GenBank prior to the start of the analysis.

There were a small number of isolates whose excessive differences between MIC tests and predictions of resistance suggested artifacts from resistance gene loss, sample swaps, testing errors, mixed cultures, or other confounding factors. We eliminated isolates where resistance calls differed from the gene-based prediction for all tested members of three or more drug classes defined as aminoglycosides, beta-lactams, lincosamides, ketolides, macrolides, phenicols, quinolones, sulfonamides, tetracyclines, and trimethoprim-sulfamethoxazole. This filter removed 38 isolates from the analyses (0.6%, Figure 1).

**Fig. 1:**
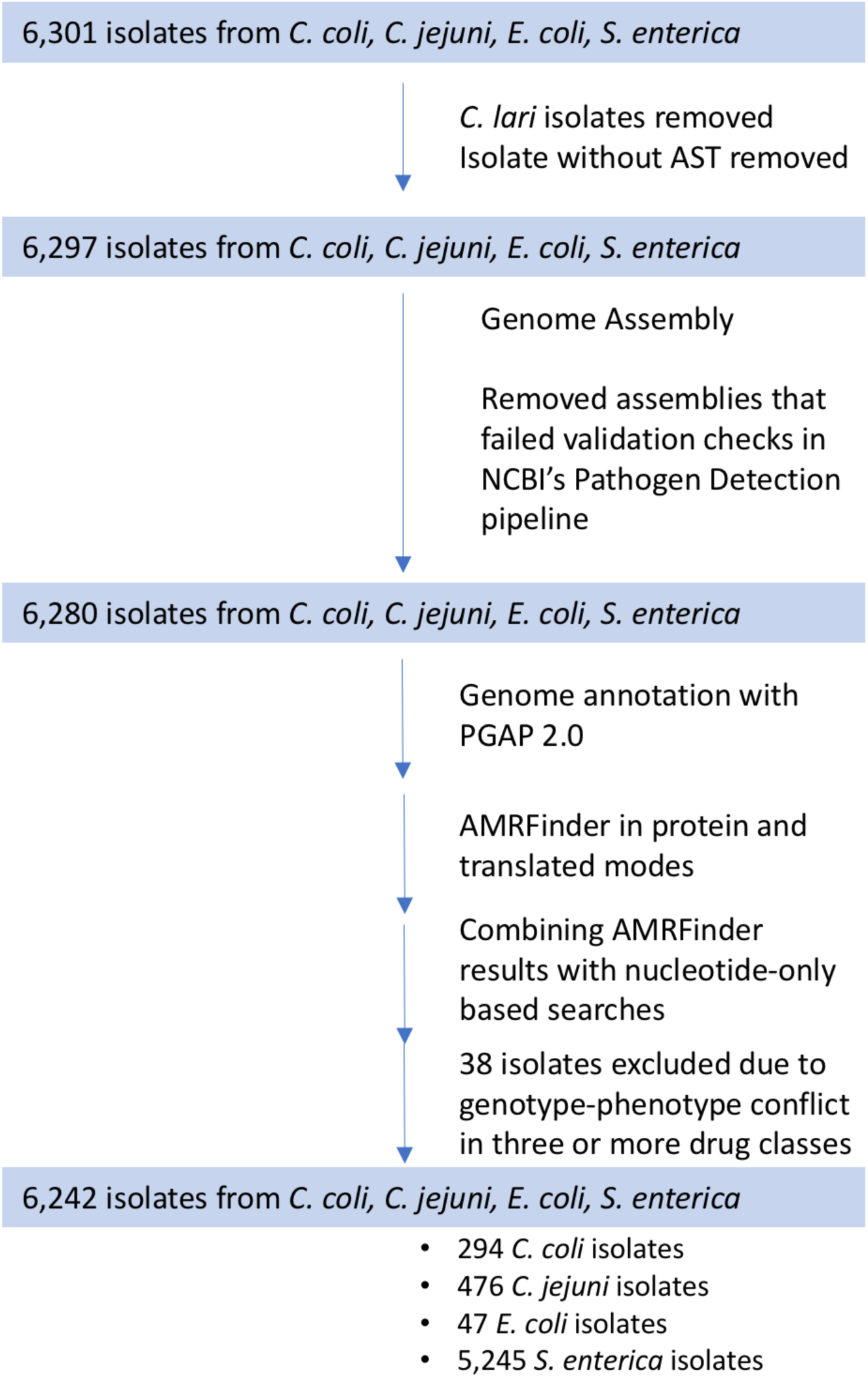
Data processing and analysis flow. Processing steps and isolate inclusion and exclusion criteria are indicated by arrows, with the number of isolates retained in each phase indicated in the colored boxes. Thirty-eight isolates were excluded if their AST phenotypes in three or more drug classes differed from predictions based on acquired AMR genes.

### Genome assembly and annotation

Illumina whole-genome shotgun reads were assembled using SPAdes v.3.5.0 using the default parameters (35). To be included in the study we required the isolate assemblies to meet the following criteria: (1) one and only one species-appropriate, full-length, *gyrA* gene; (2) < 100-Kb of the assembly in contigs covered by < 10% the genome-wide average coverage; (3) < 8-Mb in size; (4) sufficient sequence for > 20-fold genome coverage; (5) NCBI species average nucleotide identity (ANI) matched [(36) Figure 1]. To calculate coverage, reads for each isolate were aligned back to the assembly with BWA version 0.7.10-r789 using the MEM algorithm and default parameters (37). SAMtools version 1.3.1 was then used to convert alignments to read-depths for each base (38). Genomes were annotated using NCBI’s PGAP 2.0 pipeline (21,39). For 540 isolates, we used genome assemblies already deposited in GenBank (Table S1).

### Combining results

First, redundant equal-scoring hits to the same protein or identical location on the assembly were removed. Next, translated BLAST hits that overlapped over more than 75% of their length with AMRFinder-prot hits were removed as duplicates. Finally, nucleotide BLAST hits that overlapped over more than 75% of their length with either AMRFinder-prot or translated BLAST hits were removed as duplicates. 14,984 (98.19%) AMR genes were identified by the annotation-based protein AMRFinder, while 268 (1.77%) were identified by translated DNA BLAST. The remaining 7 hits (0.046%) were partial proteins identified only by nucleotide BLAST.

### Contig filtering

Reads for each isolate were aligned back to the assembly using BWA version 0.7.10-r789 using the MEM algorithm and default parameters (40). SAMtools version 1.3.1 was then used to convert alignments to read-depths for each base (38). Using this data genome-wide and per-contig average read-depths were calculated for filtering. AMR genes identified above were filtered and removed from analysis if its read-depth of the contig containing a given AMR gene was < 1/10th of the average per-base read-depth for the entire assembly.

### Identifying point mutations

Point mutations in three structural genes that confer resistance in *C. coli* and *C. jejuni* were. examined: *gyrA*, 50S ribosomal protein L22, and 23S rRNA (11). We identified putative resistance mutations by blasting the protein or nucleotide sequences against the listed accessions and predicted resistance based on the presence of the listed known resistance alleles at any of the listed offsets. The gene *gyrA* was screened (AJW58405.1 and YP_002344422.1) for the mutations T86I, T86K, T86V, D90N, D90Y, P104S, and C257T, which predict resistance to quinolones. For the 50S ribosomal protein L22 (AJW59082.1 and YP_002345068.1) we predicted resistance to macrolides due to changes at positions A84D, G86E, G86V, A88E, and A103V. The 23S rRNA (CP01115.1) was screened for those *C. jejuni* 23S mutations, A2074C, A2074G, A2074T, A2075G, and C2627A, which were expected to confer resistance to macrolides(41-43). To assess if ciprofloxacin resistance in *S. enterica* could be attributed to point mutations, we screened *gyrA* (WP_001281271.1; A67P, D72G, V73I, G81C/S/H/D, D82G/N, S83Y/F/A, D87N/G/Y/K, S97P, L98V, A119S/E/V, A131G, E139A), *gyrB* (WP_000072047.1; Y421C, R438L, S464Y/F, E466D), *parC* (WP_001281910.1; T66I, G78D, S80R/I, E84K/G), and *parE* (WP_000195318.1; M438I, E454G, S458P, V461G, H462Y, A499T, V514G, V521F) for mutations expected to confer resistance (44-47).

### Correlation of antimicrobial susceptibility phenotypes with resistance gene content

After all resistance genes were identified, isolates exhibiting phenotypic resistance were correlated with the predicted phenotype based on presence or absence of resistance genes or point mutations for each antibiotic (see Table S4 for predictions). Predicted phenotypes were scored as either resistant (R) or susceptible (S), with the presence of one or more resistance-conferring genes yielding a prediction of “R”. These were compared to the gold standard observed phenotypic results, with observed susceptibility results of intermediate (I) treated as “S”, with the exception of ciprofloxacin in *S. enterica*, for which I values were treated as resistant, since previous work has indicated that one or more resistance genes or point mutations are associated with an intermediate susceptibility phenotype (48,49).

### AMRFinder-ResFinder comparisons

AMRFinder blasts resistance gene protein sequences, either against a set of annotated proteins or a nucleotide sequence, while Resfinder uses a nucleotide database, and blasts that database against a nucleotide sequence (e.g., a bacterial genome). In addition, Resfinder reports the ‘highest-scoring’ hit, even if the underlying sequence does not support such a precise claim (e.g., calling a novel OXA allele “OXA-61”), while the hierarchical gene structure of AMRFinder will attempt to identify the appropriate gene name that does not provide an incorrect or overly precise name. To compare the output of AMRFinder to ResFinder, we first determined if these two methods called AMR genes at nearly identical coordinates on the same genome (the absolute difference in lengths could be no more than 40 bp). We used Resfinder 2.0, with the database downloaded on Nov. 15, 2017 and compared it with AMRFinder with the database locked on Feb. 2, 2017. For Resfinder, the default settings of 90% nucleotide similarity and a 60% minimum length were used. The particular version of the AMRFinder gene database used in this study can be found at ftp://ftp.ncbi.nlm.nih.gov/pathogen/Technical/AMRFinder_technical/feldgard_et_al_2018_amrdb.tar.gz. AMRFinder parameters used include a 90% nucleotide similarity and 50% minimum length for matching, and 40 disinfectant and other resistance genes were included in AMRFinder that were not in ResFinder 2.0. This allowed us to identify instances when the same gene occurred multiple times in a genome in instances where one copy was missed or misidentified by either method. We then compared gene symbols produced by each method. Where gene symbols did not agree, we assigned them to one of four categories: (1) *Synonyms* were cases where the identical protein was called by both methods, but the name differed (e.g., many aminoglycoside modifying enzymes, such as *strA* and *aph(3’’)-Ib*). (2) *Underspecified* calls occurred when the protein was 100% identical to a known, named protein, but one method did not describe it with sufficient resolution (e.g., *bla*_*TEM-1*_ is miscalled as *bla*_*TEM*_). (3) *Overspecified* calls were cases where the correct name was a less specific gene symbol, when the method provided an overspecified symbol (e.g., a novel *bla*_*TEM*_ family allele is miscalled as *bla*_*TEM-1*_). (4) *Incorrect* calls occurred when an incorrect gene symbol was ascribed to a protein (e.g., *bla*_*OXA-193*_ is miscalled as *bla*_*OXA-61*_).

### Antimicrobial susceptibility testing

Minimum inhibitory concentrations were measured using the Sensitire(tm) system and susceptibility panels designed specifically for NARMS surveillance (50). *E. coli* and *S. enterica* were tested for susceptibility to amoxicillin-clavulanic acid, ampicillin, azithromycin, cefoxitin, ceftriaxone, chloramphenicol, ciprofloxacin, trimethoprim-sulfamethoxazole, gentamicin, nalidixic acid, streptomycin, sulfisoxazole, and tetracycline; some *Salmonella* isolates were screened against amikacin, ceftiofur, kanamycin, and meropenem depending on the composition of the NARMS panel at the time of testing. *Campylobacter* spp. were screened for susceptibility to azithromycin, ciprofloxacin, clindamycin, erythromycin, florfenicol, gentamicin, nalidixic acid, telithromycin, and tetracycline.

The breakpoints used for susceptibility testing were CLSI standard breakpoints. For antibiotics that lack CLSI breakpoints, breakpoints established by the NARMS Working Group were used (Table S2, S3).

## Results

We compiled, curated, and publicly released a hierarchical database of AMR gene families, names, sequences, and HMMs with a consistent naming scheme and hierarchical structure called the Bacterial Antimicrobial Resistance Reference Gene Database (https://www.ncbi.nlm.nih.gov/pathogens/isolates#/refgene/). We also developed AMRFinder to use the AMR protein sequences, HMMs, the hierarchy of gene families and a custom rule-set to generate a report of the names, symbols, and coordinates of acquired AMR genes along with descriptions of the evidence used to identify the sequence (https://www.ncbi.nlm.nih.gov/pathogens/antimicrobial-resistance/AMRFinder/).

To verify and validate the results of the AMRFinder system, we analyzed a collection of isolates, sequenced, and susceptibility tested as part of the NARMS program. We then compared the resistance patterns predicted by AMR genes identified in the genome sequence to the results of the phenotypic susceptibility tests. We further compared the resistance gene calls made by AMRFinder to calls from the commonly used resistance gene finding tool ResFinder (22).

A total of 6,301 NARMS isolates with both phenotypes and whole-genome shotgun sequences were compiled, 59 were removed for quality reasons described above, leaving 6,242 isolates for this analysis (Figure 1). After assembly and annotation, AMRFinder was used to generate a list of 16,003 AMR gene calls, yielding 132 unique genes and alleles. Resistance predictions for the 132 genes and alleles observed in the set of 6,242 isolates were compiled from the literature (Table S4) and used to predict resistance.

### Overall consistency

For the entire set, there were 87,679 susceptibility tests performed, 98.4% (86,276) were consistent with predictions based on the resistance genotypes (acquired resistance genes, and, when tested, point mutations. Of the 13,903 tests that were predicted to be resistant, 95.5% were observed to be resistant (PPV = 0.955), while of the 73,776 tests expected to be susceptible, 99.2% were observed to be susceptible (NPV = 0.992; Table 1). 2,136 of the 6,242 isolates (34.2%) were pan-susceptible. *E. coli* isolates had the highest consistency with 99.7% (656/658) of susceptibility tests predicted by the resistance genotype. Within *S. enterica*, 98.0% of susceptibility tests were consistent with the resistance genotype, with PPV = 0.94 and NPV = 0.992 (Table 2). No resistance among *E. coli* and *S. enterica* isolates to amikacin or meropenem was observed or predicted. *C. coli* had the lowest consistency, with 96.7% of susceptibility tests consistent with the resistance genotype, with a PPV of 0.904 and an NPV of 0.982 (Table 3). 98.9% of phenotypes were accurately predicted for *C. jejuni*, with PPV = 0.971 and NPV = 0.992 (Table 4). Gentamicin and streptomycin susceptibility calls in *S. enterica* were the most common incorrect predictions, accounting for 38% of inconsistent calls (532/1,403). 17% of all isolates (1,053) had one or more inconsistent calls between genotype and phenotype.

**Table 1:** Consistency^a^ between antibiotic susceptibility phenotypes and genotype-based predictions for all NARMS isolates

|  | # resistant observations | # susceptible observations |
| --- | --- | --- |
| # predicted resistant | 13,122 | 781 |
| # predicted sensitive | 622 | 73,154 |

a Overall consistency was 98.4% of susceptibility tests performed, with a PPV = 0.955 and NPV = 0.992.

**Table 2:** *S. enterica* susceptibility and consistency^a^ with AMRFinder genotypic prediction.

| Antibiotic | # isolates susceptible<sup>b</sup> | # isolates resistant<sup>b</sup> | % consistent<sup>c</sup> | % resistant | PPV<sup>d</sup> | NPV |
| --- | --- | --- | --- | --- | --- | --- |
| amikacin | 2820 | 0 | 100.0% | 0.0% | NC | 1 |
| AMC | 4622 (7) | 718 (38) | 99.2% | 14.0% | 0.99 | 0.992 |
| ampicillin | 3734 (27) | 1620 (44) | 98.7% | 30.7% | 0.984 | 0.988 |
| azithromycin | 2592 (5) | 6 (1) | 99.8% | 0.3% | 0.545 | 0.999 |
| cefoxitin | 4686 (67) | 658 (14) | 98.5% | 12.4% | 0.908 | 0.997 |
| ceftiofur | 4093 (13) | 697 (13) | 99.5% | 14.7% | 0.982 | 0.997 |
| ceftriaxone | 4652 (8) | 744 (21) | 99.5% | 14.7% | 0.989 | 0.996 |
| CHL | 5214 (5) | 202 (4) | 99.8% | 3.8% | 0.976 | 0.999 |
| ciprofloxacin<sup>e</sup> | 5283 (14) | 114 (14) | 99.5% | 2.4% | 0.891 | 0.997 |
| cotrimoxazole | 5343 (8) | 69 (5) | 99.8% | 1.4% | 0.896 | 0.999 |
| gentamicin | 4692 (109) | 571 (53) | 97.0% | 11.5% | 0.84 | 0.989 |
| kanamycin | 3382 (23) | 412 (67) | 97.7% | 12.3% | 0.947 | 0.981 |
| meropenem | 609 | 0 | 100.0% | 0.0% | NC | 1 |
| nalidixic acid | 5294 (35) | 15 (81) | 97.9% | 1.8% | 0.3 | 0.985 |
| streptomycin | 3291 (254) | 1756 (76) | 93.9% | 33.7% | 0.877 | 0.977 |
| sulfonamide | 3763 (35) | 1572 (55) | 98.3% | 30.0% | 0.978 | 0.986 |
| tetracycline | 2558 (42) | 2776 (49) | 98.3% | 52.1% | 0.985 | 0.981 |

<sup>a</sup>Overall consistency is 98.0% of, with PPV = 0.94 and NPV = 0.992.
<sup>b</sup>The number of isolates with genotypes consistent with either phenotypic susceptibility or resistance to a given antibiotic is shown, with number of isolates with genotypes inconsistent with either susceptibility or resistance to a given antibiotic displayed in parentheses; values of zero in parentheses have been dropped for clarity.
<sup>c</sup>“% consistent” describes the percentage of isolates with a phenotype consistent with genotype.
<sup>d</sup>NC means the value cannot be calculated as there are no expected resistant isolates.
<sup>e</sup>For ciprofloxacin, # resistant included isolates with intermediate and resistant MIC results.

**Table 3:** *C. coli* susceptibility and consistency.

| Antibiotic | # isolates susceptible<sup>b</sup> | # isolates resistant<sup>b</sup> | % consistent<sup>c</sup> | % resistant | PPV<sup>d</sup> | NPV |
| --- | --- | --- | --- | --- | --- | --- |
| azithromycin | 265 | 29 | 100.0% | 9.9% | 0.763 | 1 |
| ciprofloxacin | 207 | 87 | 100.0% | 29.6% | 1 | 1 |
| clindamycin | 248 | 29 (17) | 94.2% | 15.6% | NC | 0.844 |
| erythromycin | 265 | 29 | 100.0% | 9.9% | 0.763 | 1 |
| florfenicol | 294 | 0 | 100.0% | 0.0% | NC | 1 |
| gentamicin | 288 | 6 | 100.0% | 2.0% | 1 | 1 |
| nalidixic acid | 201 (3) | 87 (3) | 98.0% | 30.6% | 1 | 0.986 |
| telithromycin | 257 (16) | 21 | 94.6% | 7.1% | 0.553 | 1 |
| tetracycline | 80 (3) | 210 (1) | 98.6% | 71.8% | 0.989 | 0.988 |

<sup>a</sup>Overall consistency was 96.7% with PPV = 0.904 and NPV = 0.982.
<sup>b</sup>The number of isolates with genotypes consistent with either susceptibility or resistance to a given antibiotic is shown, with number of isolates with genotypes inconsistent with either susceptibility or resistance to a given antibiotic displayed in parentheses. Values of zero in parentheses have been dropped for clarity.
<sup>c</sup>“% consistent” describes the percentage of isolates with a consistent phenotype. For macrolides and fluoroquinolones, consistency estimates include point mutation data.
<sup>d</sup>NC means the value can not be calculated as there are no expected resistant isolates.

**Table 4:** *C. jejuni* susceptibility and consistency^a^.

| Antibiotic | # isolates susceptible | # isolates resistant | % consistent | % resistant | PPV<sup>b</sup> | NPV |
| --- | --- | --- | --- | --- | --- | --- |
| azithromycin | 476 | 0 | 100.0% | 0.0% | 0 | 1 |
| ciprofloxacin | 386 (1) | 86 (3) | 99.2% | 18.7% | 0.989 | 0.992 |
| clindamycin | 470 | 0 (6) | 98.7% | 1.3% | NC | 0.987 |
| erythromycin | 476 | 0 | 100.0% | 0.0% | 0 | 1 |
| florfenicol | 476 | 0 | 100.0% | 0.0% | NC | 1 |
| gentamicin | 475 | 0 (1) | 99.8% | 0.2% | NC | 0.998 |
| nalidixic acid | 385 (3) | 86 (2) | 98.9% | 18.7% | 0.977 | 0.992 |
| telithromycin | 476 | 0 | 100.0% | 0.0% | 0 | 1 |
| tetracycline | 145 (4) | 325 (2) | 98.7% | 68.9% | 0.988 | 0.986 |

<sup>a</sup>Overall consistency was 98.9% with PPV = 0.971 and NPV = 0.992.
<sup>b</sup>The number of isolates with genotypes consistent with either susceptibility or resistance to a given antibiotic is shown, with number of isolates with genotypes inconsistent with either susceptibility or resistance to a given antibiotic displayed in parentheses. Values of zero in parentheses have been dropped for clarity.

### Quinolone resistance

None of the 47 *E. coli* isolates were resistant to either nalidixic acid or ciprofloxacin, nor were they predicted to be. *S. enterica* displayed high consistency for both ciprofloxacin and nalidixic acid (Table 2). When decreased susceptibility (R or I) is used as the breakpoint for ciprofloxacin (51), *S. enterica* isolates had high positive predictive values (PPV = 0.891) and high negative predictive values (NPV = 0.997). For nalidixic acid, the positive predictive value was quite low (PPV = 0.3). Thirty-five *qnr*^*+*^ isolates (71.4%) were susceptible to nalidixic acid, but they had an MIC of one doubling dilution below the nalidixic acid breakpoint of 32 μg/ml; thirteen *qnr*^*+*^ isolates were resistant to nalidixic acid; previous work indicates that qnr loci might not be very effective at conferring resistance to nalidixic acid (52). Point mutations in *gyrA* and other genes associated with ciprofloxacin were not used for the determination of nalidixic acid susceptibility, as it was unclear from some previous studies if these mutations also confer resistance to nalixidic acid (48,49). However, of the 80 isolates that had ciprofloxacin resistance mutations, 79 were resistant to nalidixic acid.

In *C. coli* and *C. jejuni*, fluoroquinolone resistance was associated with point mutations, not acquired genes (Tables 3, 4). Based on previous reports (11), we examined the relationship between *gyrA* mutations previously determined to confer fluoroquinolone resistance and fluoroquinolone resistant isolates among these *Campylobacter* spp. isolates. All but two fluoroquinolone resistant and no fluoroquinolone susceptible *C. coli* isolates possessed a GyrA T86I mutation (Table S5). In *C. jejuni*, 84/85 isolates with GyrA T86I mutations were resistant to ciprofloxacin, and 83/85 were resistant to nalidixic acid; three *C. jejuni* isolates without known fluoroquinolone resistance mutations were resistant to both fluoroquinolones; no unique mutations were correlated with these three isolates.

Thus in *S. enterica*, presence of *qnr* genes or QRDR mutations conferred either resistance or decreased susceptibility to ciprofloxacin, while in *Campylobacter* spp. *gyrA* mutations conferred resistance.

### Macrolides and lincosamides

Only six of eleven *S. enterica* isolates predicted to be azithromycin resistant were resistant. These six resistant isolates carried *mph(A)*; however, one azithromycin susceptible isolate also carried *mph(A)*. The other four susceptible isolates carried either the *ere(A)* or *abc-f* resistance genes; these isolates did not have elevated MICs near the top end of the susceptible range.

All *C. jejuni* were susceptible to azithromycin, erythromycin, and telithromycin, with only six *C. jejuni* displaying resistance to clindamycin (Table 4). None of the clindamycin resistant *C. jejuni* isolates had any known resistance mutations or unique mutations suggesting novel resistance mutations in either 23S or the 50S/L22 subunit (Table S5). Macrolide resistance was far more common in *C. coli* (Table 3), with most resistant isolates possessing a A2075G mutation in 23S (Table S6), as has been observed previously (11).

### Decreased amoxicillin-clavulanic acid susceptibility in S. enterica

As expected, we observed that 718 out of 725 *S. enterica* isolates (99.0%) with one or more *bla*_CMY_*-*family genes were resistant to amoxicillin-clavulanic acid. As observed previously (51), other beta-lactamases conferred decreased or intermediate susceptibility to amoxicillin-clavulanic acid (Fig. 3). 92.6% of isolates that carried a *bla*_PSE_*/bla*_CARB_ family beta-lactamase (a novel *bla*_CARB_ allele or *bla*_CARB-2_) displayed intermediate susceptibility to amoxicillin-clavulanic acid, while over half of those isolates with a *bla*_HER_ family beta-lactamase displayed intermediate susceptibility to amoxicillin-clavulanic acid, with the remainder having a MIC of 8 μg/ml, which is the highest MIC categorized as susceptible. *bla*_TEM_ isolates had a similar pattern, with nearly half displaying intermediate susceptibility.

**Fig. 2 a, b:**
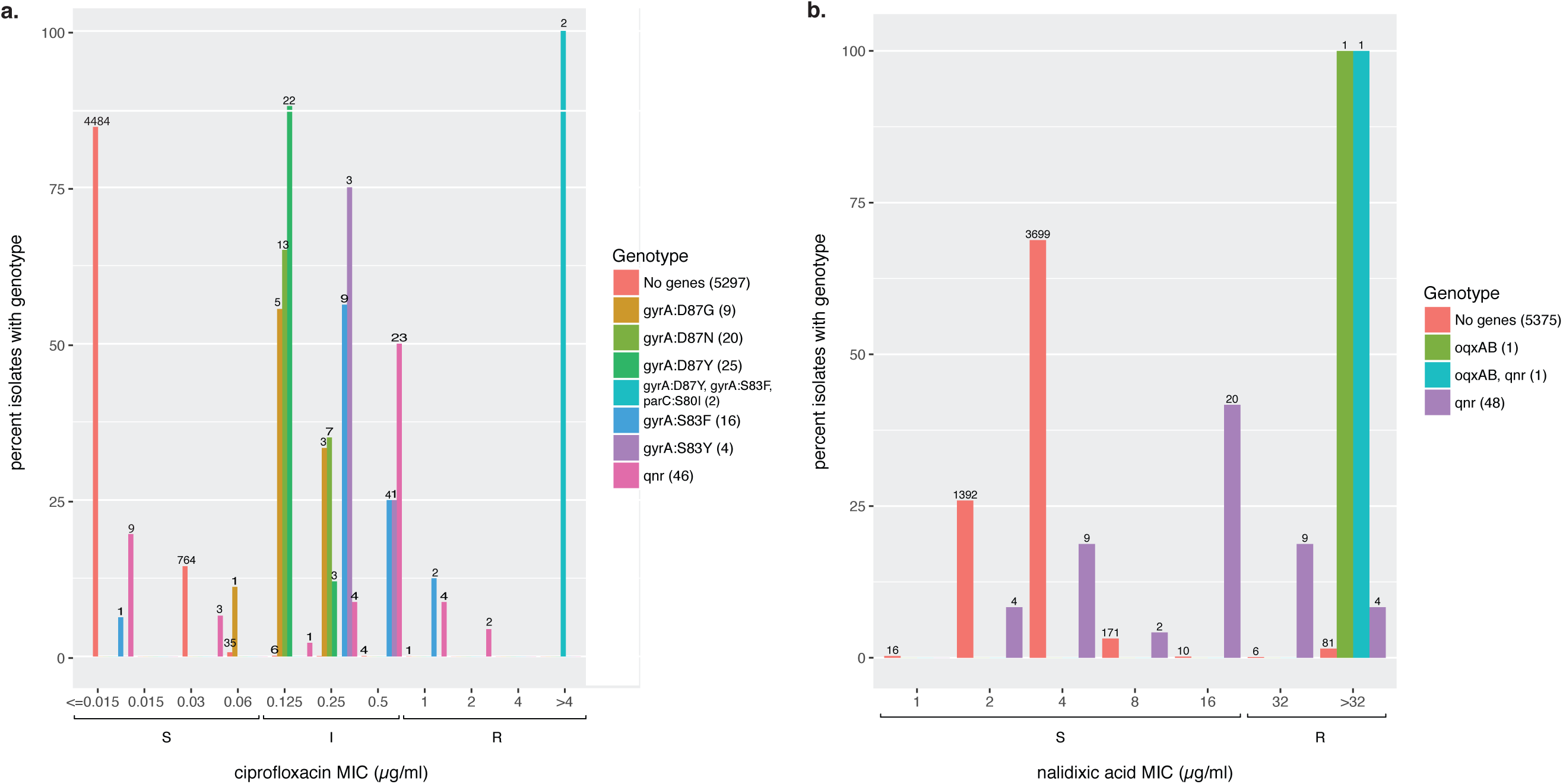
Qnr loci affect ciprofloxacin (a) and nalidixic acid (b) MICs in *S. enterica*. Columns on the x-axis correspond to observed MIC values; brackets below indicate the SIR values for those MICs. On the y-axis, colored bars indicate the percentage of isolates sharing the same genotype with a given MIC value. Numbers above each column indicate the number of isolates observed with that MIC and genotypes. In the side legend, the number in parentheses is the number of isolates with the corresponding genotype. “No genes” are those isolates lacking any predicted fluoroquinolone resistance genes. *oqxAB* indicates the presence of these fluorquinolone resistance genes in an isolate. “qnr” indicates the presence of one of the following Qnr family genes: QnrB2, QnrB19, QnrB77, QnrS1, QnrS2, or an unassigned QnrB family allele. “oqxAB, qnr” indicates an OqxAB, QnrB19 genotype. Point mutations are indicated by the gene in which they occurred, followed by the site and changed residues.

**Fig. 3:**
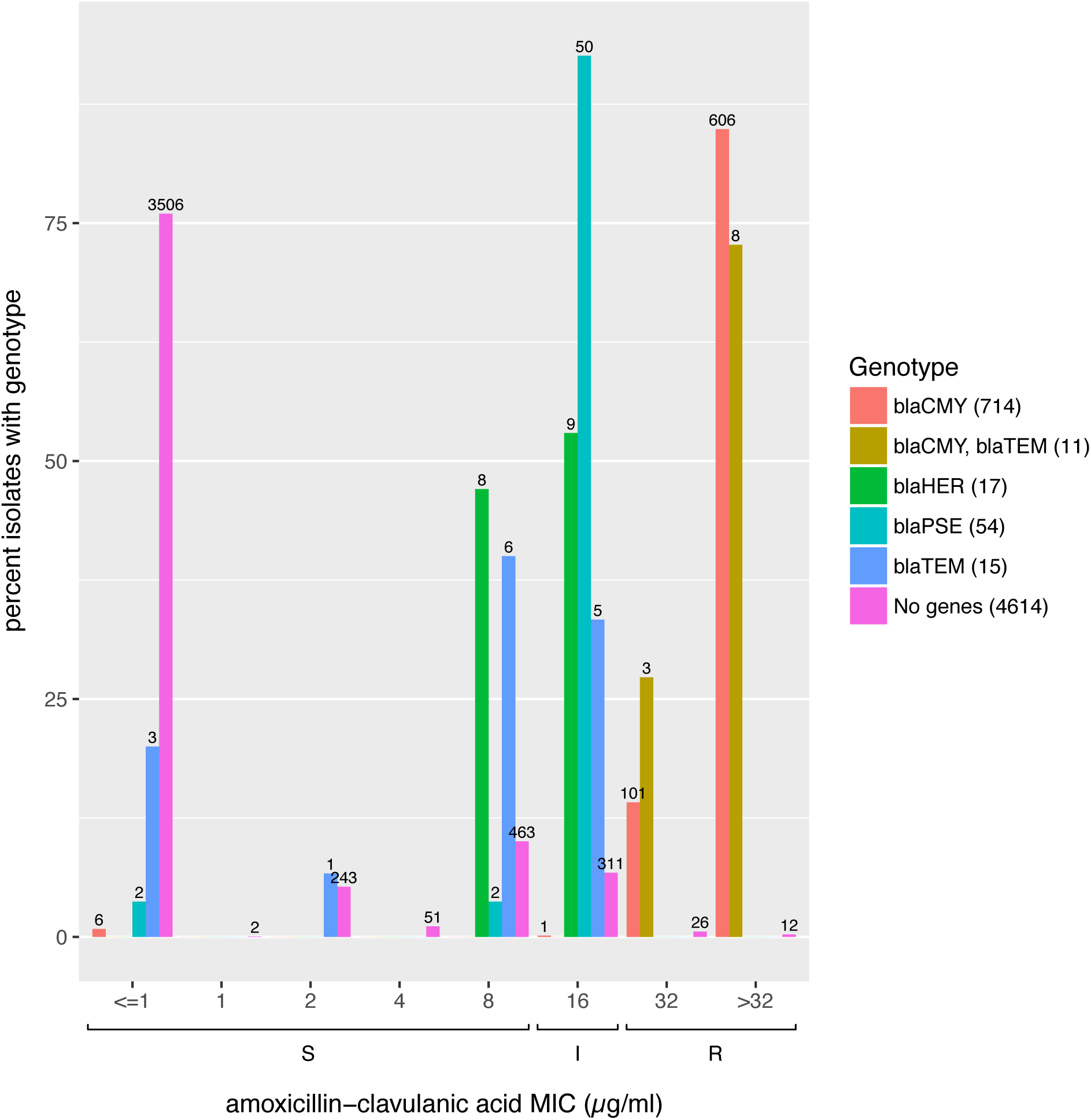
Unexpected beta-lactamases confer decreased susceptibility to amoxicillin-clavulanic acid in *S. enterica*. X-and y-axis as above. Allelic variants within a beta-lactamase family are grouped together under the family name; an isolate can have multiple alleles belonging to the same family. “blaPSE” family beta-lactamases are either CARB-2 or unassigned CARB alleles. “blaCMY” family beta-lactamases were either novel *bla*_*CMY*_ alleles or the CMY-2 allele. “blaHER” indicates either the HER-3 allele or a novel HER-family allele. “blaTEM” indicates either a novel TEM allele, or TEM-1 “No genes” indicates those isolates lacking beta-lactamases.

### Aminoglycoside susceptibility in Salmonella

Overall, the presence or absence of acquired gentamicin and kanamycin resistance genes was a good predictor of susceptibility phenotypes (Table 2). Of the 2,820 *Salmonella* that were tested for susceptibility to amikacin, none were resistant, nor were they predicted to be resistant. However, we noticed that several reported gentamicin and kanamycin resistance genes conferred decreased susceptibility to gentamicin and kanamycin even if the MICs were not high enough to qualify as resistant (Fig 4a, b). The majority of *aac(3)-IV*^*+*^ isolates (36/47) and 26% of *ant(2″)-Ia*^*+*^ isolates displayed intermediate susceptibility to gentamicin. Many *aac(6’)-Ib*^*+*^ isolates were susceptible to gentamicin, but the MICs of these isolates were higher than isolates lacking known resistance genes. While *aac(6’)-Ib* family enzymes, other than *aac(6’)-Ib4*, do not confer resistance to gentamicin, they are known to confer resistance to some of the individual components of gentamicin, such as gentamicin C1a and C2, and thus these genes might decrease susceptibility to gentamicin (53). While most kanamycin resistance genes were associated with phenotypic resistance, 13% of *ant(2″)-Ia*^*+*^ isolates had intermediate susceptibility.

**Fig. 4 a, b:**
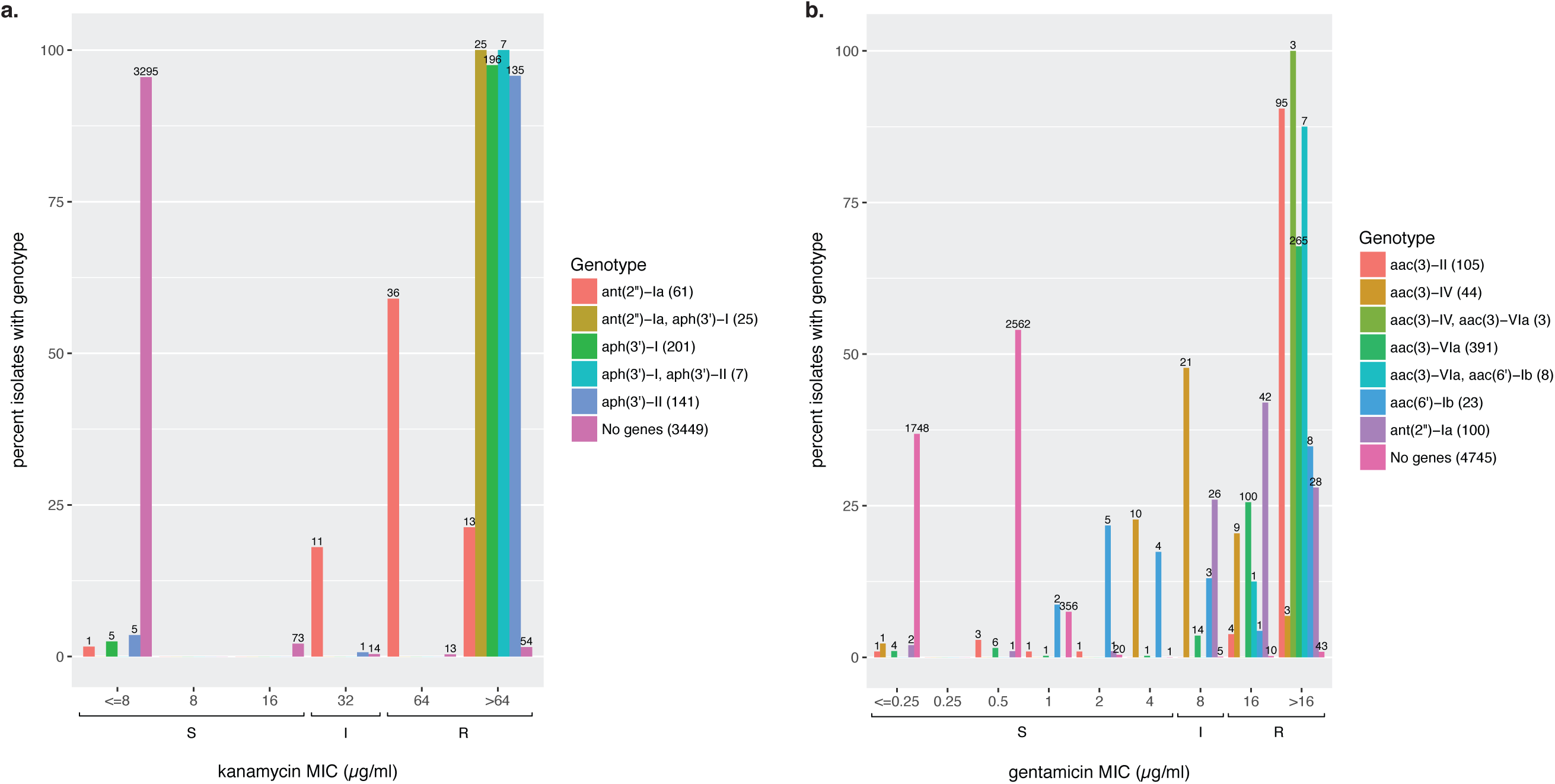
Gentamicin and kanamycin resistance in *S. enterica*. Format as described for Figure 2 except aminoglycoside modifying genes are grouped together by family. “No genes” are those isolates lacking any predicted gentamicin and kanamycin resistance genes respectively.

As noted previously, streptomycin susceptibility calls accounted for a large fraction of the inconsistent calls, with many such isolates containing putative streptomycin genes. There were no obvious direct relationships between particular resistance genes and streptomycin susceptibility (see Table S7). We examined whether partial genes (defined as 50%-90% of the closest reference protein length) affected susceptibility calls. Partial genes only accounted for 6.4% of streptomycin discrepancies, suggesting this observation is not due to potential non-functional genes. While the mechanism of discordance between streptomycin resistance genes and susceptibility is unclear, this relationship has been observed in multiple surveys of Enterobacteriaceae (14, 16, 51, 54, 55). [also see S. 8]

### AMRFinder-ResFinder comparison

ResFinder is a widely used AMR determinant detection program(22). To assess the relative accuracy of AMRFinder we compared the gene symbol calls at similar positions in the two tools. As described in Methods, discrepant gene symbol calls were classified into four different categories: synonyms, overspecification (e.g., calling a novel or partial *bla*_*TEM*_ allele as *bla*_*TEM-1*_), underspecification (e.g., calling an actual *bla*_*TEM-1*_ allele a *bla*_*TEM*_ -family allele), and miscalls (e.g., mislabeling a full-length, 100% identical sequence as a different, known full-length sequence).

Overall, out of 14,023 AMR genes identified by both AMRFinder and ResFinder there were 1,229 gene symbol discrepancies (Tables 5, S8). These discrepancies could be mapped to 42 gene symbols, out of a total of 132 unique AMRFinder gene symbol calls. ResFinder misidentified 247 genes with an exact match to a known AMR gene or allele (e.g., misidentifying *blaOXA-193* as *blaOXA-61*), and over-specified the gene symbol in 977 cases, representing 18 misidentified gene symbols and 21 overspecified gene symbols out of the set of 132 unique AMR gene symbols. In five cases, AMRFinder underspecified the gene symbol, representing three underspecifications out of the set of 132 unique AMR protein symbols.

**Table 5:** Discrepancies by category observed in gene symbol calls by AMRFinder and ResFinder 2.0 from 2017.

| Error type<sup>a</sup> | AMRFinder | ResFinder |
| --- | --- | --- |
| Misclassification | 0 | 247 |
| Underspecification | 5 | 0 |
| Overspecification | 0 | 977 |

<sup>a</sup>Synonyms are not included in this table as they do not represent miscalls by either system.

The ResFinder misclassifications resulted from either the absence of the matching sequence in the ResFinder database used in this study or a lack of correspondence between the closest nucleotide hit and actual observed sequence. For example, 32 *aac(6’)-Ib* family genes, including 22 known, 100% identity *aac(6’)-Ib4* sequences, were miscalled as *aac(6’)-Ib-cr*. The gene *aac(6’)-Ib-cr* contributes to decreased fluoroquinolone susceptibility and confers amikacin and tobramycin resistance, while *aac(6’)-Ib4* does not confer resistance or decreased susceptibility to amikacin, ciprofloxacin, or tobramycin. We would note that none of the sixteen *S. enterica aac(6’)-Ib4*^*+*^ isolates that also were tested for susceptibility to amikacin were resistant to amikacin, supporting the AMRFinder call of *aac(6’)-Ib4*. In 977 instances, ResFinder overspecified the gene symbol as it calls the closest hit as the correct gene symbol. Many of these were novel, unnamed allelic variants of beta-lactamase families (n = 699; Table S8), and Resfinder reported the closest hit (e.g., *blaOXA-61* when a novel *blaOXA* sequence was observed).

We also examined the loci that were missed by either ResFinder or AMRFinder. ResFinder did not find 1,147 AMR loci that AMRFinder identified (Table 6). Most of the missed loci (81.2%) belonged to drug or disinfectant classes that ResFinder does not cover, bleomycin and quarternary ammonium compounds. Bleomycin resistance is included in the AMRFinder database and is highly associated with the clinically relevant NDM family carbapenemases (56), although both databases do look directly for NDM genes, while *qac* enzymes can be linked to multiple resistance genes (57). The next largest class belonged to AMR genes that were not represented in the ResFinder database (8.8%). The default setting length of 60% of the reference sequence also resulted in 111 missed calls. Of 66 genes not found by ResFinder that could be assessed by susceptibility data (out of the total of 111), 53 genes were consistent with the susceptibility data (associated with a resistant phenotype), while thirteen were not.

**Table 6:** Unique proteins found by AMRFinder

| <u>Explanation</u> | <u>N (tot=1,147)</u> | <u>%</u> |
| --- | --- | --- |
| Drug class not in ResFinder | 931 | 81.2 |
| Proteins below thresholds<sup>a</sup> | 111 | 9.7 |
| Gene not found in ResFinder | 101 | 8.8 |
| Translation/frameshift errors<sup>b</sup> | 4 | 0.3 |

<sup>a</sup>In ten cases, ResFinder was unable to detect these as the nucleotide sequence was too divergent from any sequence found in the database. In 101 instances, there was no gene in the ResFinder database with >90% DNA sequence similarity to the predicted genes.
<sup>b</sup>Frameshifts led to early stop codons, resulting in stop codon positions that differed by more than 40bp between the two methods.

AMRFinder missed 16 loci that ResFinder found. In all 16 cases, these were frameshifts or in-frame stop codons that resulted in a translated protein that either was not identified at all or had a stop codon position that differed from the ResFinder stop position by more than 40 bp. Of the three loci that AMRFinder missed that were assessed phenotypically, all of which were frameshifts, two were resistant in spite of the apparent frameshift, while one was susceptible. There were also 21 instances of an *aph(6)-I* like gene that was divergent from AMR genes in either the ResFinder or the AMRFinder protein database. Due to this divergence, the two systems identified proteins that differed in length and thus had divergent start and stop sites, and were therefore called as misses.

## Discussion

We developed and populated a highly curated database with hierarchical structure for AMR proteins, with tuned cutoffs and associated hierarchical names. AMRFinder uses this AMR protein database, HMMs, a hierarchy of AMR protein families, and a custom rule-set to identify AMR genes. In addition, AMRFinder reports the evidence used to make the determination users can evaluate its strength and their confidence in the calls.

We observed high consistency between the presence of acquired AMR determinants and resistance phenotypes. We would note, however, that, as part of our sample consisted of isolates that were resistant to one or more antibiotics, our choice of isolates might overestimate the overall PPV, while underestimating the NPV. Incorporating mutational resistance also increased PPV and decreased NPV for certain drugs, especially fluoroquinolones and macrolides, as resistance to these drugs was predominantly mutational and not due to acquired AMR genes. The *E. coli* sample was small (n = 47), and most *E. coli* isolates were susceptible to most antibiotics, leading to very high consistency. In *S. enterica*, discrepancies in aminoglycoside resistance and fluoroquinolone resistance typically arose from acquired resistance genes conferring intermediate MICs or MICs at the high end of the susceptible range. As other studies in foodborne pathogens have demonstrated (51,54), clinical breakpoints, while obviously critical for appropriate treatment, do not always correspond to the presence or absence of resistance genes.

Beta-lactam resistance in *S. enterica* showed high correlation between resistance phenotypes and genotypes overall. Elevated MICs and intermediate susceptibility amoxicillin-clavulanic acid phenotypes in *S. enterica* were associated with the presence of beta-lactamases other than *bla*_*CMY*_. NCBI’s Pathogen Detection system (http://ncbi.nlm.nih.gov/pathogens), as part of a collaboration with the FDA GenomeTrakr (58), CDC PulseNet (59), and USDA-FSIS, routinely clusters genomes by sequence similarity, including the isolates described in this report, to support outbreak and traceback investigations of clonal isolates. We determined that these isolates belong to different SNP clusters, and so it does not appear that this pattern stems from chance sampling of a single clone with an unknown resistance mechanism, though we cannot rule out an unknown, common mechanism of decreased susceptibility. One possible explanation why *bla*_PSE_ family, *bla*_HER_, and *bla*_TEM_ beta-lactamase carrying isolates would display this phenotypic difference could be that these beta-lactamases are overproduced in the presence of amoxicillin-clavulanic acid; overexpression of *bla*_TEM-1_ in *E. coli* confers amoxicillin-clavulanic acid resistance(60). Alternatively, changes in permeability or efflux could lower the intracellular concentration of either the drug or the inhibitor, conferring intermediate or decreased susceptibility.

As found in previous studies, resistance to macrolides and quinolones in these *C. coli* and *C. jejuni* (11) is largely due to point mutations. When we screened for point mutations in *gyrA* and 23S, we were able to predict phenotypes with extremely high accuracy. This highlights the importance of point mutations in determining resistance phenotypes. Future editions of AMRFinder will incorporate point mutation information for *Campylobacter, E. coli*, and *S. enterica*.

Comparing AMRFinder to ResFinder revealed the importance of annotation and of a comprehensive AMR reference gene database. Protein length variation, when working with AMR proteins, can yield false conclusions. For any AMR gene detection system, incomplete or incorrect databases can lead to AMR gene identification errors.

We also found that there were instances where the highest scoring ResFinder hit was either incorrect due to absence of a sequence specific enough to make the correct call or to a reference nucleotide sequence that was divergent from the correct sequence. One case was the *aac(6’)* family aminoglycoside modifying enzyme. Slight nucleotide changes that result in protein differences can result in the gain or loss of fluoroquinolone and aminoglycoside resistance (61). We also observed miscalls of QnrB alleles (quinolone resistance) and OXA-61 family beta-lactamases due to the closest nucleotide hit not corresponding to the correct protein hit. AMRFinder, by having a nested hierarchical classification of AMR proteins into families, is able to appropriately name novel AMR genes, which can avoid imputing incorrect function by overspecifing the gene name. Without a clear interpretation of what similarity, but not complete identity, to known AMR genes means, using a ‘highest scoring hit’ approach can lead to false conclusions regarding AMR gene content.

Although allele miscalls might appear to be minor, and in many cases might not affect susceptibility patterns, there are cases where these differences have profound effects on the predicted resistance phenotype. As mentioned above, very minor differences in aminoglycoside modifying enzymes can result in significant differences in susceptibility. Recent work with KPC family beta-lactamases has revealed that a subset of alleles, including *bla*_KPC-8_, are not only resistant to carbapenemases, but also ceftazidime-tazobactam (62). *bla*_KPC-8_ was first described in 2008 before ceftazidime-tazobactam existed as a treatment option. In some circumstances, accurate identification down to the allele level is crucial to characterizing the relationship between resistance genotype and phenotype. Comparisons in this study used older versions of both the AMRFinder and Resfinder databases out of necessity, as both systems are continuously improving their databases. Since we locked down the databases for both systems, as of Sept. 1, 2018, the Resfinder database has grown from 2,254 nucleotide sequences to 3,307 (a 35% increase), and the AMRFinder database has increased by 17%, from 3,921 protein sequences to 4,579. These improvements should increase the accuracy of both systems.

Note that reliability of WGS-based methods is dependent on the accuracy of the underlying WGS data. Low-level contamination or poor-quality sequence data can lead to inaccurate assessments; this is a particular problem with ‘greedy’ assemblers that will assemble very low coverage regions. Consensus assemblers run the risk that nearly identical orthologous genes or low-level sequencing contamination might yield an incorrect sequence. Low-quality assemblies can also result in partial genes, making assessment of resistance genes challenging. To increase the accuracy and reliability of AMR gene identification, NCBI is developing an assembler that emphasizes base accuracy, increasing the reliability of allele identification (63).

In analyzing these data, we also encountered several issues. There are two competing, partially overlapping aminoglycoside modifying enzyme nomenclature systems. This makes constructing reference gene databases, as well as validating them, extremely difficult. We also discovered that, in developing the genotype-phenotype matrix, there are many alleles and genes that either have not been characterized phenotypically at all, or only against a subset of antibiotics. This was a particular problem with the beta-lactamases, where in some cases alleles were characterized phenotypically before the advent of currently used drugs. In addition, some genes are described very broadly. Terms such as ‘cephalosporin-hydrolyzing’ or ‘aminoglycoside-modifying’ do not aid accurate prediction. While these terms can be useful when confronted with a novel allele or gene, in that they avoid making unwarranted statements about phenotype, we would encourage more phenotypic assessment of novel and existing genes using well-standardized methods and quality control, such as the CLSI or EUCAST standards, to guide WGS-based methods and increase our basic understanding of AMR. It would also help to have more phenotypic data publicly available and linked to existing genome sequences (https://www.ncbi.nlm.nih.gov/biosample/docs/antibiogram/).

In AMRFinder, we have adopted a protein-focused approach, as opposed to a nucleotide-oriented approach, for several reasons. First, protein annotation and similarity comparisons against both reference proteins and using HMMs with appropriate cutoffs can aid in determining if the gene is functional, whereas a nucleotide approach can miss nonsense mutations. Second, the protein sequence encodes the AMR function. Even single amino acid changes can significantly alter resistance phenotypes, and this variation should be explicitly captured. Third, there can be discordance between nucleotide and protein sequences, leading to the mis-assignment of alleles, and thus potentially to incorrect prediction of AMR phenotypes. Note, however, that there can be upstream mutations that interfere with gene expression, and that these types of mutations are not being reported by AMRFinder. For example, *bla*_KPC_ alleles in the context of different *Tn4401* variants are expressed at different levels (64,65). Even when we used both nucleotide and protein approaches, and removed isolates that had genotype-phenotype discrepancies among three or more drug classes, we still observed that 17% of isolates had one or more discrepancies between the resistance genotype and the observed antibiogram. Even with high consistency for individual tests, isolates tested on multiple drugs will likely have one or more discrepancies as a simple statistical property. For example, 21% of isolates tested against twelve antibiotics with a consistency of 98% would have one or more errors (assuming an equal consistency rate for each antibiotic). Further technical refinements will be needed to lower the per-isolate discrepancy further, if clinical prediction is a primary goal.

The tool we have described, AMRFinder, uses a combined protein BLAST and HMM approach. BLAST can identify complete or near matches to known genes. HMMs based on curated data, on the other hand, can identify putative resistance genes that fall below arbitrary BLAST thresholds, enabling the recognition of novel resistance genes. By integrating both of these methods, we are able to assign the most specific functional name possible to the AMR protein (66).

While AMRFinder is a powerful tool for identifying acquired resistance genes, our *Campylobacter* results highlight the importance of assessing the role of point mutations. To better understand the context in which AMR genes occur, NCBI is also developing a biocide and metal resistance database to screen for genes linked to resistance to those compounds. The latest AMRFinder software, source code, and databases are publicly available at https://www.ncbi.nlm.nih.gov/pathogens/antimicrobial-resistance/AMRFinder/. While this study examined foodborne pathogens, NCBI’s Pathogen Detection system, which facilitates the analysis of food-borne and clinical isolates to aid outbreak and traceback investigations, uses AMRFinder to identify AMR genes from over 200,000 clinical and environmental bacterial isolates (https://www.ncbi.nlm.nih.gov/pathogens/), enabling the rapid identification of isolates with important AMR-related genotypes.

## Acknowledgements

This work was supported by the Intramural Research Program of the National Institutes of Health, National Library of Medicine.

## Disclaimer

The views expressed in this article are those of the authors and do not necessarily reflect the official policy of the Department of Health and Human Services, the U.S. Food and Drug Administration, the Centers for Disease Control and Prevention, or the U.S. Government. Mention of trade names or commercial products in this publication is solely for the purpose of providing specific information and does not imply recommendation or endorsement by the Food and Drug Administration.

