## Supplemental Tables 2-6 for "Using the NCBI AMRFinder Tool to Determine Antimicrobial Resistance Genotype-Phenotype Correlations Within a Collection of NARMS Isolates"

Supplemental Material:

| | *C. jejuni* | *C. jejuni* | *C. coli* | *C. coli* | Standard[a] |
|---|---|---|---|---|---|
| antibiotic | S | R | S | R | |
| gentamicin | <=2 | >=4 | <=2 | >=4 | NARMS |
| telithromycin | <=4 | >=8 | <=4 | >=8 | NARMS |
| clindamycin | <=0.5 | >=1 | <=1 | >=2 | NARMS |
| azithromycin | <=0.25 | >=0.5 | <=0.5 | >=1 | NARMS |
| erythromycin | <=4 | >=8 | <=8 | >=16 | CLSI |
| chloramphenicol | <=16 | >=32 | <=16 | >=32 | NARMS |
| ciprofloxacin | <=0.5 | >=1 | <=0.5 | >=1 | CLSI |
| nalidixic acid | <=16 | >=32 | <=16 | >=32 | NARMS |
| doxycycline | <=0.5 | >=1 | <=1 | >=2 | NARMS |
| tetracycline | <=1 | >=2 | <=2 | >=4 | CLSI |

Table S2: *Campylobacter* sp. breakpoints used in this study.
[a]Standard refers to whether the breakpoint was established by CLSI or, if there is no CLSI breakpoint, by NARMS. Units are μg/ml.

| antibiotic | S | I | R | Standard[a] |
|---|---|---|---|---|
| gentamicin | <= 4 | 8 | >=16 | CLSI |
| streptomycin | <=32 | N/A | >=64 | NARMS |
| amoxicillin-clavulanic acid | <=8/4 | 16/8 | >=32/16 | CLSI |
| cefoxitin | <=8 | 16 | >=32 | CLSI |
| ceftiofur | <=2 | 4 | >=8 | CLSI |
| ceftriaxone | <=1 | 2 | >=4 | CLSI |
| sulfamethoxazole/sulfisoxazole | <=256 | N/A | >=512 | CLSI |
| co-trimoxazole | <=2/38 | N/A | >=4/76 | CLSI |
| azithromycin | <=16 | N/A | >=32 | NARMS |
| ampicillin | <=8 | 16 | >=32 | CLSI |
| chloramphenicol | <=8 | 16 | >=32 | CLSI[b] |
| ciprofloxacin | <=0.06 | 0.12-0.5 | >=1 | CLSI[c] |
| nalidixic acid | <=16 | N/A | >=32 | CLSI |
| tetracycline | <= 4 | 8 | >=16 | CLSI |
| amikacin | <=16 | 32 | >=64 | CLSI |

Table S3: *E. coli* and *S. enterica* breakpoints.

[a]Standard refers to whether the breakpoint was established by CLSI or, if there is no CLSI breakpoint, by NARMS. Units are μg/ml.
[b]For E. coli isolates, the NARMS convention of using Salmonella breakpoints was followed.
[c]For S. enterica, isolates were scored as resistant to ciprofloxacin if they were intermediately susceptible (“I”).

| Gene as determined by AMRFinder | Predicted antibiotic resistance |
|---|---|
| *aac(3)* | gentamicin |
| *aac(3)-I* | gentamicin |
| *aac(3)-Id* | gentamicin |
| *aac(3)-II* | gentamicin |
| *aac(3)-IId* | gentamicin |
| *aac(3)-IV* | gentamicin |
| *aac(3)-VIa* | gentamicin |
| *aac(6')-Ib* | amikacin |
| *aac(6')-Ib4* | gentamicin |
| *aac(6')-Ie* | amikacin, gentamicin |
| *aac(6')-IIc* | gentamicin |
| *aadA* | streptomycin |
| *aadA1* | streptomycin |
| *aadA12* | streptomycin |
| *aadA13* | streptomycin |
| *aadA15* | streptomycin |
| *aadA2* | streptomycin |
| *aadA21* | streptomycin |
| *aadA22* | streptomycin |
| *aadA25* | streptomycin |
| *aadA29* | streptomycin |
| *aadA4* | streptomycin |
| *aadA5* | streptomycin |
| *aadA6* | streptomycin |
| *aadA7* | streptomycin |
| *aadA8* | streptomycin |
| *aadE* | streptomycin |

| | |
|---|---|
| *abc-f* | azithromycin |
| *ampC* | amoxicillin-clavulanic acid, ampicillin |
| *ant(2″)-Ia* | gentamicin, kanamycin |
| *ant(3″)* | streptomycin |
| *ant(6)* | streptomycin |
| *ant(6)-Ia* | streptomycin |
| *aph(2″)-Ig* | amikacin, gentamicin |
| *aph(2″)-IIIa* | amikacin, gentamicin |
| *aph(3″)* | streptomycin |
| *aph(3″)-Ib* | streptomycin |
| *aph(3′)-I* | kanamycin |
| *aph(3′)-Ia* | kanamycin |
| *aph(3′)-Id* | kanamycin |
| *aph(3′)-II* | kanamycin |
| *aph(3′)-IIa* | kanamycin |
| *aph(3′)-IIIa* | amikacin, kanamycin |
| *aph(3′)-VIIa* | kanamycin |
| *aph(6)-I* | streptomycin |
| *aph(6)-Ic* | streptomycin |
| *aph(6)-Id* | streptomycin |
| *armA* | amikacin, gentamicin |
| $bla_{CARB-2}$ | ampicillin |
| $bla_{CMY}$ | amoxicillin-clavulanic acid, ampicillin, cefoxitin, ceftiofur, ceftriaxone |
| $bla_{CMY-2}$ | amoxicillin-clavulanic acid, ampicillin, cefoxitin, ceftiofur, ceftriaxone |

| | |
|---|---|
| *bla*$_{CMY\text{-}5}$ | amoxicillin-clavulanic acid, ampicillin, cefoxitin, ceftiofur, ceftriaxone |
| *bla*$_{CMY\text{-}61}$ | amoxicillin-clavulanic acid, ampicillin, cefoxitin, ceftiofur, ceftriaxone |
| *bla*$_{CMY\text{-}7}$ | amoxicillin-clavulanic acid, ampicillin, cefoxitin, ceftiofur, ceftriaxone |
| *bla*$_{CTX\text{-}M}$ | ampicillin, ceftriaxone |
| *bla*$_{CTX\text{-}M\text{-}1}$ | ampicillin, ceftriaxone |
| *bla*$_{CTX\text{-}M\text{-}24}$ | ampicillin, ceftriaxone |
| *bla*$_{CTX\text{-}M\text{-}65}$ | ampicillin, ceftriaxone |
| *bla*$_{HER}$ | ampicillin |
| *bla*$_{HER\text{-}1}$ | ampicillin |
| *bla*$_{HER\text{-}3}$ | ampicillin |
| *bla*$_{LAT\text{-}1}$ | amoxicillin-clavulanic acid, ampicillin, cefoxitin, ceftiofur, ceftriaxone |
| *bla*$_{OXA}$ | ampicillin |
| *bla*$_{OXA\text{-}1}$ | ampicillin |
| *bla*$_{OXA\text{-}184}$ | ampicillin, amoxicillin-clavulanic acid, cefoxitin, ceftriaxone, meropenem |

| | |
|---|---|
| *bla*$_{OXA-193}$ | ampicillin, amoxicillin-clavulanic acid, cefoxitin, ceftriaxone, meropenem |
| *bla*$_{OXA-2}$ | ampicillin |
| *bla*$_{OXA-449}$ | ampicillin |
| *bla*$_{OXA-450}$ | ampicillin |
| *bla*$_{OXA-460}$ | ampicillin |
| *bla*$_{OXA-461}$ | ampicillin |
| *bla*$_{OXA-489}$ | ampicillin |
| *bla*$_{OXA-493}$ | ampicillin |
| *bla*$_{OXA-61}$ | ampicillin |
| *bla*$_{SHV-12}$ | ampicillin, ceftriaxone |
| *bla*$_{SHV-2A}$ | ampicillin, ceftriaxone |
| *bla*$_{TEM}$ | ampicillin |
| *bla*$_{TEM-1}$ | ampicillin |
| *bla*$_{TEM-116}$ | ampicillin |
| *bla*$_{TEM-135}$ | ampicillin |
| *catA1* | chloramphenicol |
| *catA2* | chloramphenicol |
| *catB3* | chloramphenicol |
| *cmlA* | chloramphenicol |

| | |
|---|---|
| *cmlA1* | chloramphenicol |
| *cmlA5* | chloramphenicol |
| *cmlA6* | chloramphenicol |
| *dfr7* | trimethoprim-sulfamethoxazole* |
| *dfrA1* | trimethoprim-sulfamethoxazole* |
| *dfrA12* | trimethoprim-sulfamethoxazole* |
| *dfrA14* | trimethoprim-sulfamethoxazole* |
| *dfrA15* | trimethoprim-sulfamethoxazole* |
| *dfrA17* | trimethoprim-sulfamethoxazole* |
| *dfrA19* | trimethoprim-sulfamethoxazole* |
| *dfrA5* | trimethoprim-sulfamethoxazole* |
| *dfrB* | trimethoprim-sulfamethoxazole* |
| *dfrI* | trimethoprim-sulfamethoxazole* |
| *ere(A)* | azithromycin, erythromycin |
| *erm(42)* | azithromycin, clindamycin, erythromycin |
| *floR* | chloramphenicol, florfenicol |
| *lnu(G)* | clindamycin |
| *mef(B)* | azithromycin, erythromycin |
| *mph(A)* | azithromycin, erythromycin |
| *mph(E)* | azithromycin, erythromycin |
| *msr(E)* | azithromycin, erythromycin |
| *oqxA* | ciprofloxacin, nalidixic acid |
| *oqxB* | ciprofloxacin, nalidixic acid |
| *qnrA1* | ciprofloxacin, nalidixic acid |
| *qnrB* | ciprofloxacin, nalidixic acid |
| *qnrB19* | ciprofloxacin, nalidixic acid |
| *qnrB2* | ciprofloxacin, nalidixic acid |
| *qnrB77* | ciprofloxacin, nalidixic acid |
| *qnrS1* | ciprofloxacin, nalidixic acid |

| | |
|---|---|
| *qnrS2* | ciprofloxacin, nalidixic acid |
| *sul1* | sulfamethoxazole, sulfisoxazole, trimethoprim-sulfamethoxazole* |
| *sul1delta* | sulfamethoxazolesulfisoxazole, trimethoprim-sulfamethoxazole* |
| *sul2* | sulfamethoxazole, sulfisoxazole, trimethoprim-sulfamethoxazole* |
| *sul3* | sulfamethoxazole, sulfisoxazole, trimethoprim-sulfamethoxazole* |
| *tet* | tetracycline |
| *tet(32)* | tetracycline |
| *tet(A)* | tetracycline |
| *tet(B)* | tetracycline |
| *tet(C)* | tetracycline |
| *tet(D)* | tetracycline |
| *tet(G)* | tetracycline |
| *tet(M-W-O-S)* | tetracycline |
| *tet(M)* | tetracycline |
| *tet(O)* | tetracycline |
| *tetA* | tetracycline |

Table S4: Predicted susceptibility based on acquired resistance.
*trimethoprim-sulfamethoxazole requires both *dfr*-family and *sul* family genes.

| Species | drug | mutation[a] | n | # resistant (%) | # sensitive (%) |
|---|---|---|---|---|---|
| *C. coli* | ciprofloxacin | GyrA:T86I | 87 | 87 (100%) | 0 |
| *C. coli* | ciprofloxacin | none observed | 207 | 0 | 207 (100%) |
| *C. jejuni* | ciprofloxacin | GyrA:T86I | 85 | 84 (99%) | 1 (1%) |
| *C. jejuni* | ciprofloxacin | GyrA:T86K | 1 | 1 (100%) | 0 |
| *C. jejuni* | ciprofloxacin | GyrA:T86V | 1 | 1 (100%) | 0 |
| *C. jejuni* | ciprofloxacin | none observed | 389 | 3 (1%) | 386 (99%) |
| *C. coli* | nalidixic acid | GyrA:T86I | 87 | 87 (100%) | 0 |
| *C. coli* | nalidixic acid | none observed | 207 | 3 (2%) | 204 (98%) |
| *C. jejuni* | nalidixic acid | GyrA:T86I | 85 | 83 (98%) | 2 (2%) |
| *C. jejuni* | nalidixic acid | GyrA:T86K | 1 | 1 (100%) | 0 |
| *C. jejuni* | nalidixic acid | GyrA:T86V | 1 | 1 (100%) | 0 |
| *C. jejuni* | nalidixic acid | none observed | 389 | 3 (1%) | 386 (99%) |

Table S5: Known fluoroquinolone resistance mutations are highly correlated with fluoroquinolone resistance in *Campylobacter* spp. [a]Mutations refer to the position in GyrA (AJW58405.1 for *C. coli*; YP_002344422.1 for *C. jejuni*). “None observed” describes those isolates lacking any previously described fluoroquinolone resistance mutations.

| Species | drug | mutation[a] | n | # resistant (%) | # sensitive (%) |
|---|---|---|---|---|---|
| *C. coli* | azithromycin | 23S:A2075G | 29 | 29 (100%) | 0 |
| *C. coli* | azithromycin | no mutation | 265 | 0 | 265 (100%) |
| *C. coli* | clindamycin | 23S:A2075G | 29 | 29 (100%) | 0 |
| *C. coli* | clindamycin | no mutation | 265 | 17 (6%) | 248 (94%) |
| *C. coli* | erythromycin | 23S:A2075G | 29 | 29 (100%) | 0 |
| *C. coli* | erythromycin | no mutation | 265 | 0 | 265 (100%) |
| *C. coli* | telithromycin | 23S:A2075G | 29 | 21/29 (72%) | 8/29 (28%) |
| *C. coli* | telithromycin | no mutation | 265 | 0 | 265 (100%) |

Table S6: Known macrolide resistance mutations are highly correlated with fluoroquinolone resistance in *C. coli*.
[a]Mutations refer to the position in *C. coli* 23S (CP011015.1, pos. 39330-42399, VC76_00160). "No mutations" describes those isolates lacking any previously described macrolide resistance mutations.
